# A Single Mechanosensitive Channel Protects *Francisella tularensis* subsp. *holarctica* from Hypoosmotic Shock and Promotes Survival in the Aquatic Environment

**DOI:** 10.1101/198184

**Authors:** David R. Williamson, Kalyan K. Dewan, Tanmay Patel, Catherine M. Wastella, Gang Ning, Girish S. Kirimanjeswara

## Abstract

*Francisella tularensis* subspecies *holarctica* is found throughout the northern hemisphere and causes the disease tularemia in humans and animals. An aquatic cycle has been described for this subspecies, which has caused water-borne outbreaks of tularemia in at least 10 countries. In this study, we sought to identify mechanosensitive channel(s) required for the bacterium to survive the transition from mammalian hosts to freshwater, which is likely essential for transmission of the bacterium between susceptible hosts. A single <u>m</u>echano<u>s</u>ensitive <u>c</u>hannel MscS (FTL_1753), among the smallest members of the mechanosensitive channel superfamily, was found to protect subsp. *holarctia* from hypoosmotic shock. Deletion of this channel did not affect virulence within the mammalian host, however *mscS* was required to survive the transition from the host niche to fresh water. Deletion of *mscS* did not alter the sensitivity of *F. tularensis* subspecies *holarctica* to detergents, H_2_O_2_, or antibiotics, suggesting that the role of MscS is specific to protection from hypoosmotic shock. Interestingly, deletion of *mscS* also led to reduced average cell size without altering gross cell morphology. The small mechanosensitive channel identified and characterized in this study likely contributes to the transmission of tularemia between hosts by allowing the bacterium to survive the transition from mammalian hosts to fresh water.

## Importance

Contamination of fresh water by *Francisella tularensis* subspecies *holarctica* has resulted in a number of outbreaks of tularemia. Invariably, contamination originates from the carcasses or excreta of infected animals, and thus involves an abrupt osmotic shock as the bacteria enter fresh water. How *F. tularensis* survives this drastic change in osmolarity has not been clear, but here we report that a single mechanosensitive channel protects the bacterium from osmotic downshock. This channel is functional despite being notably smaller (165 a.a.) than those found in model organisms (~280 a.a.). These findings extend our understanding of the aquatic cycle and ecological persistence *of F. tularensis*, with further implications for mechanosensitive channel biology.

## Introduction

*Francisella tularensis* is a gram-negative bacterium responsible for the disease tularemia in humans and a wide range of animal species. Two subspecies of *F. tularensis* are of clinical significance for humans, ssp. *tularensis* and *holarctica* (1). Subspecies *tularensis* is considered more virulent, and is primarily confined to Northern America. Subspecies *holarctica* on the other hand is broadly distributed throughout the northern hemisphere, and has also been found in Australia (2). The manifestations of tularemia vary depending upon the route of exposure. The ulceroglandular and glandular forms of tularemia are more frequent and are associated with exposure via arthropod bites or direct contact with infected animals, leading to localized lymphadenopathy with or without the formation of an ulcer at the inoculation site (1, 3, 4). Another common manifestation is oropharyngeal tularemia, where ingestion of contaminated water leads to pharyngitis and swelling of the cervical lymph nodes (1). Rare presentations include oculoglandular tularemia, involving conjunctivitis resulting from inoculation of the eye; as well as typhoidal and pneumonic tularemia, which are both severe systemic diseases resulting from inhalation of *F. tularensis* (pneumonic) or with no obvious route of exposure (typhoidal)(3).

Tularemia is considered a zoonotic disease that is rarely, if ever, transmitted by human-to-human contact (3). Hence, understanding how the bacterium persists and spreads in the environment is important. While long-term reservoirs and transmission cycles of *F. tularensis* are not well understood (5), the role of fresh water is of particular interest in the ecology of the broadly distributed *F. tularensis* subsp. *holarctica*, for which an aquatic cycle has been described (1). Many large outbreaks of tularemia are of the oropharyngeal form, and are linked to direct ingestion of water contaminated with *F. tularensis* subsp*. holarctica*. Outbreaks of tularemia from contaminated drinking water have been reported in at least 10 countries, totaling thousands of cases. Such outbreaks have been reported in Turkey (6–8), Sweden (9), Germany 10), Norway 11–13), Russia (14, 15), Bulgaria (16, 17), Kosovo (18, 19), The Czech Republic(20), Italy (21, 22) and the Republic of Georgia (23). Isolated cases of tularemia associated with infection from fresh water have also been described in France (24, 25) and the USA (26). In many of these cases, *F. tularensis* was detected by PCR (10–13, 20) or cultured (8, 9, 15–17, 23, 26) from water of the affected area.

As the presence of viable *F. tularensis* subsp. *holarctica* in fresh water is directly responsible for many naturally occurring cases of oropharyngeal tularemia, it is important to understand how the bacterium adapts and survives in water. *F. tularensis* subsp. *holarctica* has been shown to survive in otherwise sterile fresh water for 7-40 days (27, 28). Persistence may be enhanced by the presence of amoebae or ciliates (29, 30), but the bacterium is not able to persist in fresh water indefinitely (27, 28), and is often only seasonally recovered from natural bodies of water (7, 31). This suggests that *F. tularensis* is introduced to bodies of water shortly before outbreaks, and indeed almost all cases of water contamination are attributed to the carcasses or excreta of infected animals including hares, rodents, voles, lemmings, muskrats, and beavers (6–8, 10–12, 16, 17, 19, 22, 26, 31–33). The shift from an animal to a fresh water niche involves an abrupt decrease in extracellular osmolarity, producing a hypoosmotic shock (or ‘downshock’).

Downshock results in an rapid influx of water into the cytoplasm, increasing tension on the membrane and potentially leading to lysis of the cell (34). To prevent this outcome, bacteria rely on mechanosensitive channels, which open in response to physical stretch of the membrane and jettison osmolytes (35). By doing so, mechanosensitive channels relieve pressure, restore isotonicity, and allow survival of hypoosmotic shock (36). In this study, we sought to identify the mechanosensitive channel(s) that protect *F. tularensis* subsp. *holarctica* from hypoosmotic shock, allowing it to survive the transition from a host to fresh water. We bioinformatically identified prospective mechanosensitive channels in subsp. *holarctica*, deleted candidate genes, and characterized the resulting mutants. We show that a single Mechanosensitive Ion Channel protein (MscS) is required for surviving the transition from mammalian cells to fresh water. Notably, the mechanosensitive channel gene identified herein is among the smallest members of the mechanosensitive channel superfamily. Bacteria lacking this channel display a specific defect in survival of osmotic shock, -while sensitivities to detergents, H_2_O_2_ and antibiotics were not affected. We also demonstrate that deletion of this gene results in a decrease in cell size, but does not affect the bacterium’s ability to replicate in macrophages or cause disease in a mouse model of tularemia. Together, these data indicate that MscS is critical for survival of *F. tularensis* in fresh water environments, and plays a major role in the bacterium’s ecological persistence.

## Results

### F. tularensis Live Vaccine Strain mechanosensitive channels closely resemble those found in clinical and environmental strains

We first sought to compare putative mechanosensitive channels in various *F. tularensis* subsp. *holarctica* strains, in order to determine whether the Live Vaccine Strain (LVS) was an accurate representation of other members of the subspecies. We compared LVS to a human clinical isolate (FSC200), an isolate from a beaver (OSU18) and an isolate from drinking water (R13-38). Our search yielded four loci predicted to code for mechanosensitive channels in each genome (FTL_1753, FTL_0945, FTL_1588, and FTL_1209 in LVS). Two genes (FTL_1753 and FTL_0945) were identical in all four strains of subsp. *holarctica* (Fig. S1 & S2). When we performed our initial bioinformatic comparison, FTL_1588 was annotated as a pseudogene in RefSeq (8-12-2015 annotation with NCBI software revision 3.0). Nevertheless, our analysis of FTL_1588 revealed that it was identical between LVS and FSC200 (Fig. S3). However, in OSU18, a small deletion at the beginning of the open reading frame, a potential membrane localization sequence, was observed (Fig. S3). A gap in the WGS assembly of R13-38 precluded this strain from comparison. The final hit was annotated as a pseudogene (FTL_1209) due to frameshift in all four strains. Taken together, these data show that the Live Vaccine Strain codes for mechanosensitive channels and is an accurate representation of clinical and environmental subsp. *holarctica* strains.

### A single mechanosensitive channel protects F. tularensis subsp. holarctica from hypoosmotic shock

We next sought to determine which of the predicted mechanosensitive channels conferred protection from hypoosmotic shock/downshock. Because FTL_1753 and FTL_0945 were found in all four strains of *F. tularensis* subspecies *holarctica*, marker-less, in-frame deletions of these two genes were generated in the Live Vaccine Strain (Fig. S4). A hypoosmotic shock of 300mM had little impact on the viability of WT or ΔFTL_0945 strains, but killed >90% of ΔFTL_1753 cells, indicating this gene is required for bacterial survival following an osmotic downshock (Fig. 1A). To determine the magnitude of osmotic shock required to kill ΔFTL_1753 in media, we measured survival after graded downshocks of 100, 200 or 300mM. The downshock of 300mM killed the vast majority ofΔFTL_1753 cells as before; 200mM downshock caused a moderate but significant loss of viability, and a 100mM downshock did not result in any loss of viability (Fig. 1B). These data for ΔFTL_1753 are similar to prior observations for *E. coli* lacking two mechanosensitive channel genes, *mscS* and *mscL* (36). *Trans*-complementation with an expression plasmid for FTL_1753 completely restored the ability of ΔFTL_1753 to survive all three graded downshocks (Fig. 1B). Finally, we tested whether the observed phenotype was relevant to surviving the transition from a mammalian host to fresh water, by equilibrating bacteria in PBS then diluting them into natural lake water or double distilled water. While ΔFTL_1753 was unable to survive in lake water and distilled water, wild-type, ΔFTL_0945, and *trans*-complemented strain of ΔFTL_1753 were able to survive in both conditions (Fig. 1C). These data demonstrate that FTL_1753, but not FTL_0945, encodes a functional mechanosensitive channel that is required by *F. tuluarensis* subsp. *holarctica* to survive hypoosmotic shock, including that encountered when the bacterium transitions from a mammalian host to fresh water.

**Figure 1:**
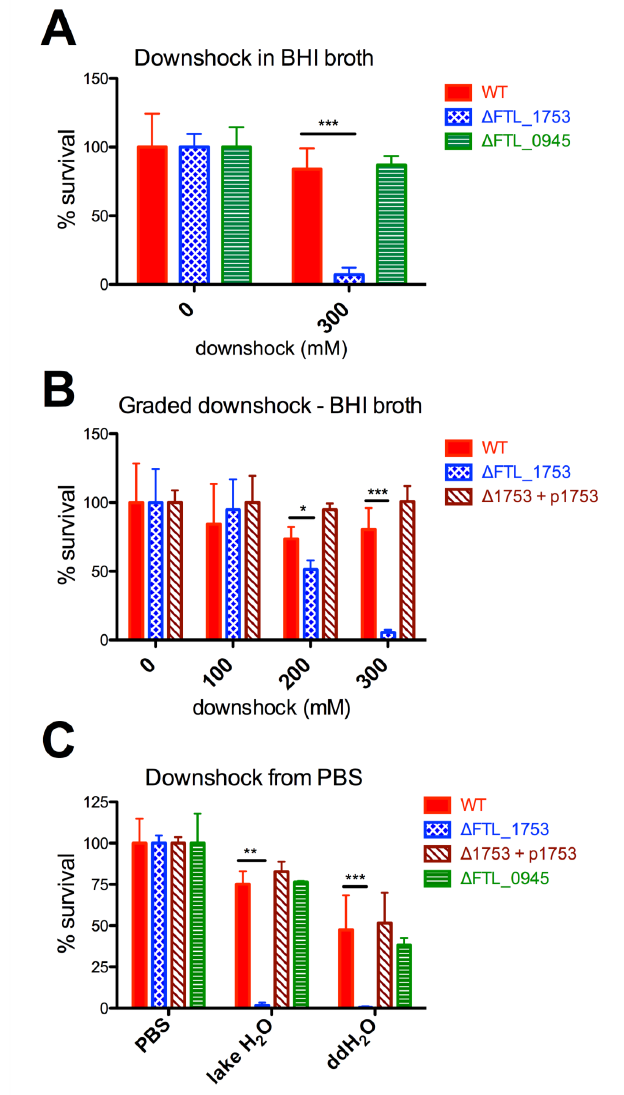
FTL_1753 is required for *F. tularensis* subsp*. holarctica* to survive hypoosmotic shock. **A-B)** Survival data from downshock experiments performed in BHI broth. Bacteria were grown in BHI + 300mM NaCl, then diluted in BHI +300, +200, +100 or +0 mM NaCl, producing hypoosmotic shocks of 0, 100, 200 and 300mM, respectively. **C)** survival data for PBS-equilibrated bacteria diluted into water retrieved from a freshwater lake or double-distilled H_2_O. Statistical differences were determined by one-way ANOVA with Dunnett’s post test, *p<0.05,**p<0.01,** i*p<0.001. Each experiment was performed a minimum of three times.

### ΔFTL_1753 bacteria are not more sensitive to detergents, H2O2 or antibiotics

To establish whether deletion of FTL_1753 or FTL_0945 resulted in more generalized vulnerability to stressors, we tested these strains for sensitivity to three detergents and H_2_O_2._ The three detergents included an anionic detergent (SDS), a cationic detergent (CTAB), and a nonionic detergent (Triton-X100). In contrast to the large differences observed in survival of downshock, all strains displayed a similar sensitivity profile to these stressors (Fig. 2). As mechanosensitive channels have been associated with antibiotic sensitivity in some contexts (41–43), we determined the minimum inhibitory concentrations (MICs) of 9 antibiotics for these strains. The knockout strains displayed identical sensitivity as the wild-type, with the exception of ΔFTL_1753 appearing slightly more resistant to kanamycin (Table S1). These data support a specific role for MscS in survival of hypoosmotic shock, rather than a more general role in resistance of physical or chemical stressors.

**Figure 2:**
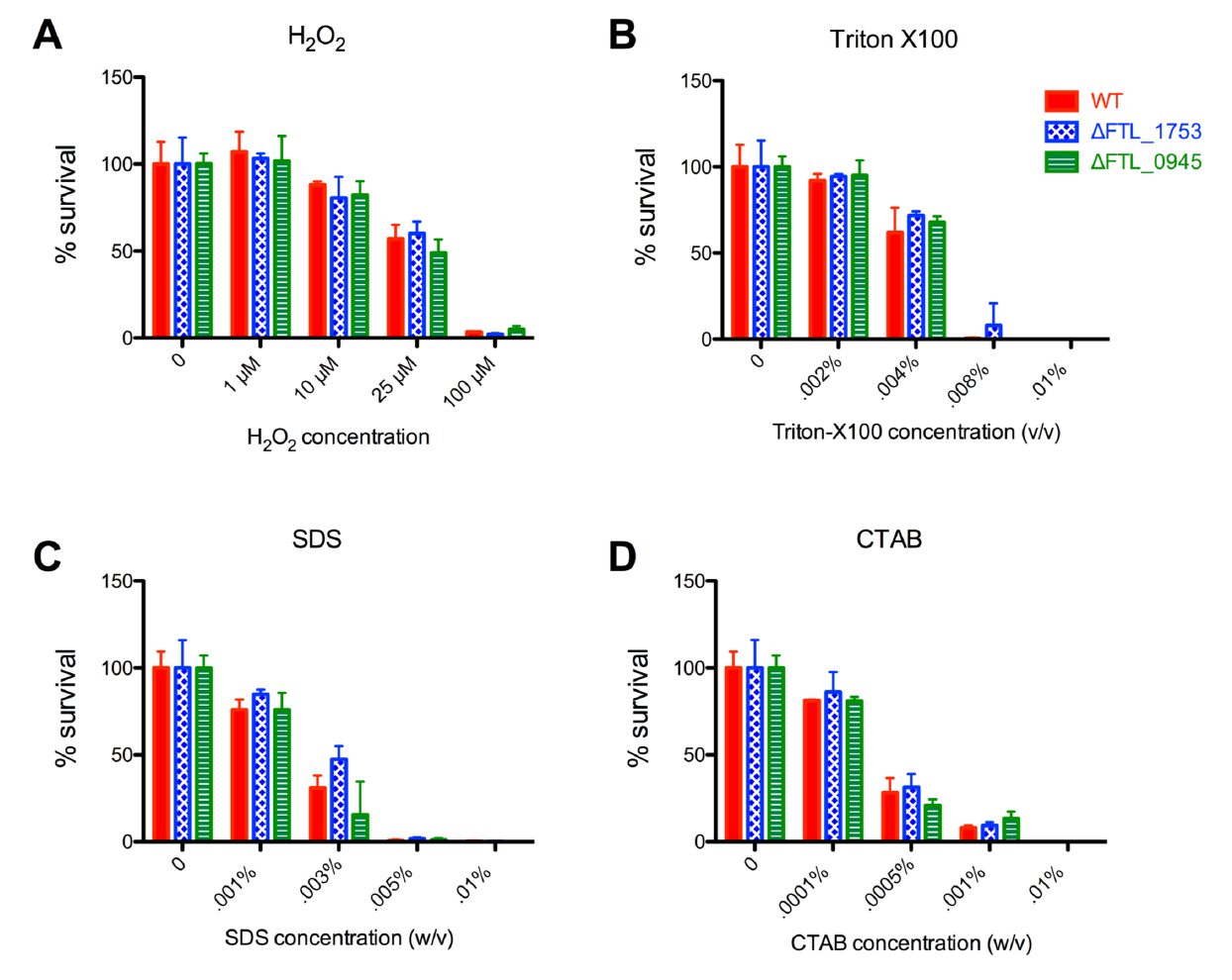
Sensitivity profiles of wild-type, ΔFTL_1753 and ΔFTL_0945 to detergents and H_2_O_2_. Early log phase bacteria were exposed to the indicated concentrations of detergents/H_2_O_2_ in media for one hour before dilution and plating to enumerate viable CFUs. Representative results shown from a minimum of four repeats. No statistically significant differences were found by one-way ANOVA with Dunnett’s post-test.

### MscS does not contribute to virulence in the host niche, but is required to survive the transition from that niche to fresh water

We next sought to determine the roles of FTL_1753 and FTL_0945 in the context of the mammalian host niche. As *F. tularensis* is an intracellular pathogen that predominantly targets macrophages, we tested the ability of these strains to proliferate in macrophages using gentamicin protection assays. Similar intra-macrophage growth was observed for all strains, suggesting that these genes do not contribute to *F. tularensis* replication within host cells (Fig. 3A). For the most part, mammalian tissues are maintained at similar osmolarities. However, some large osmotic gradients are found in mammals, most notably in the Kidneys and Liver. To determine whether FTL_1753 or FTL_0945 contribute to the pathogenesis of *F. tularensis* in the mammalian host, we tested these mutants in a mouse model of pneumonic tularemia. Mice infected with all strains displayed similar survival profiles, suggesting that these genes do not contribute to virulence within a host (Fig. 3B). Finally, we tested whether FTL_1753 is required for surviving the transition from a mammalian cell niche to fresh water, using modified gentamicin protection assay. After eliminating extracellular bacteria, infected macrophages were lysed with either detergent in PBS, or with ddH_2_O. Similar numbers of viable bacteria were recovered using the PBS-detergent lysis buffer. In contrast, FTL_1753 was required for bacteria to survive macrophage lysis with ddH_2_O (Fig. 3C). Collectively, these data suggest that MscS is not required for virulence within the mammalian host niche, but is required to survive the transition from this niche to fresh water.

**Figure 3:**
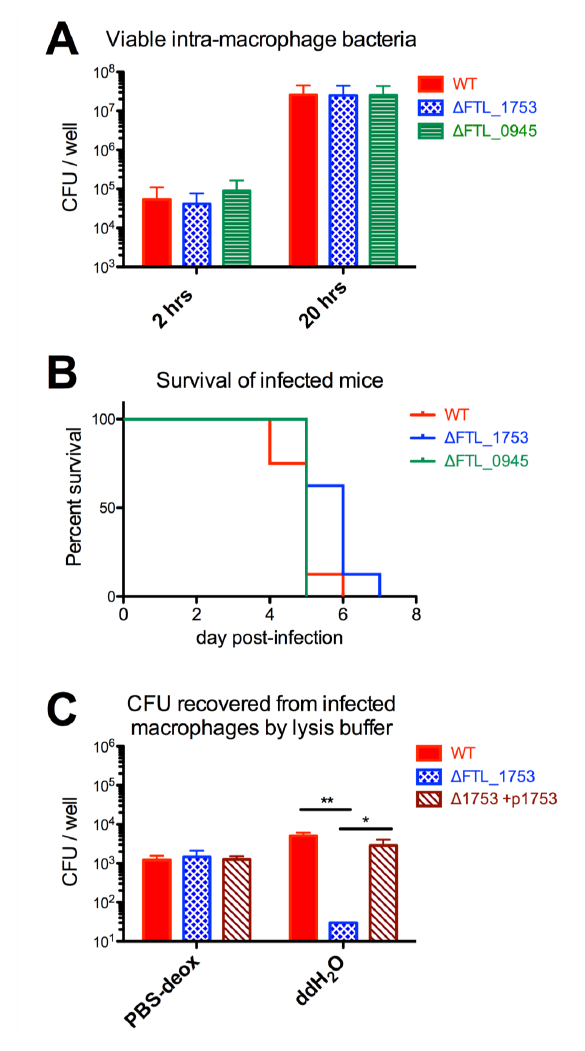
FTL_1753 does not contribute to virulence in the host niche, but is required to survive the transition from that niche to fresh water. **A)** Viable CFUs of indicated *F. tularensis* strains recovered from Raw 264.7 macrophages in a gentamicin protection assay at 2 and 20 hours post-infection, pooled data from three repeats. **B)** Survival curves of C57BL/6J mice (n=8 per group) infected with 10^4^ CFU of the indicated strains of *F. tularensis* LVS. **C)** Viable CFUs of indicated *F. tularensis* strains recovered from infected RAW 264.7 macrophages based on lysis buffer used. PBS-deox: 0.1% (w/v) sodium deoxycholate in PBS. One-way ANOVA with Tukey’s post-test, *p<0.05, ** p<0.01.

### ΔFTL_1753 cells are smaller in average size compared to the wild-type

As there are reports of mechanosensitive channels affecting cellular morphology (41), and we observed that ΔFTL_1753 often formed smaller colonies than the wild-type, we examined wild-type and ΔFTL_1753 cells using scanning electron microscopy. ΔFTL_1753 cells were of smaller average size than the wild-type; 2D outline traces revealed significantly smaller areas and perimeters compared to the wild-type (Fig. 4A-F). Although smaller in average size, ΔFTL_1753 cells had a similar overall morphology to the wild-type, and there was no difference in the aspect ratio of fitted elliptical axes (Fig. 4G). These data show that the absence of FTL_1753 leads to smaller average bacterial cell size, but does not lead to other gross morphological changes compared to wild-type *F. tularensis* LVS.

**Figure 4:**
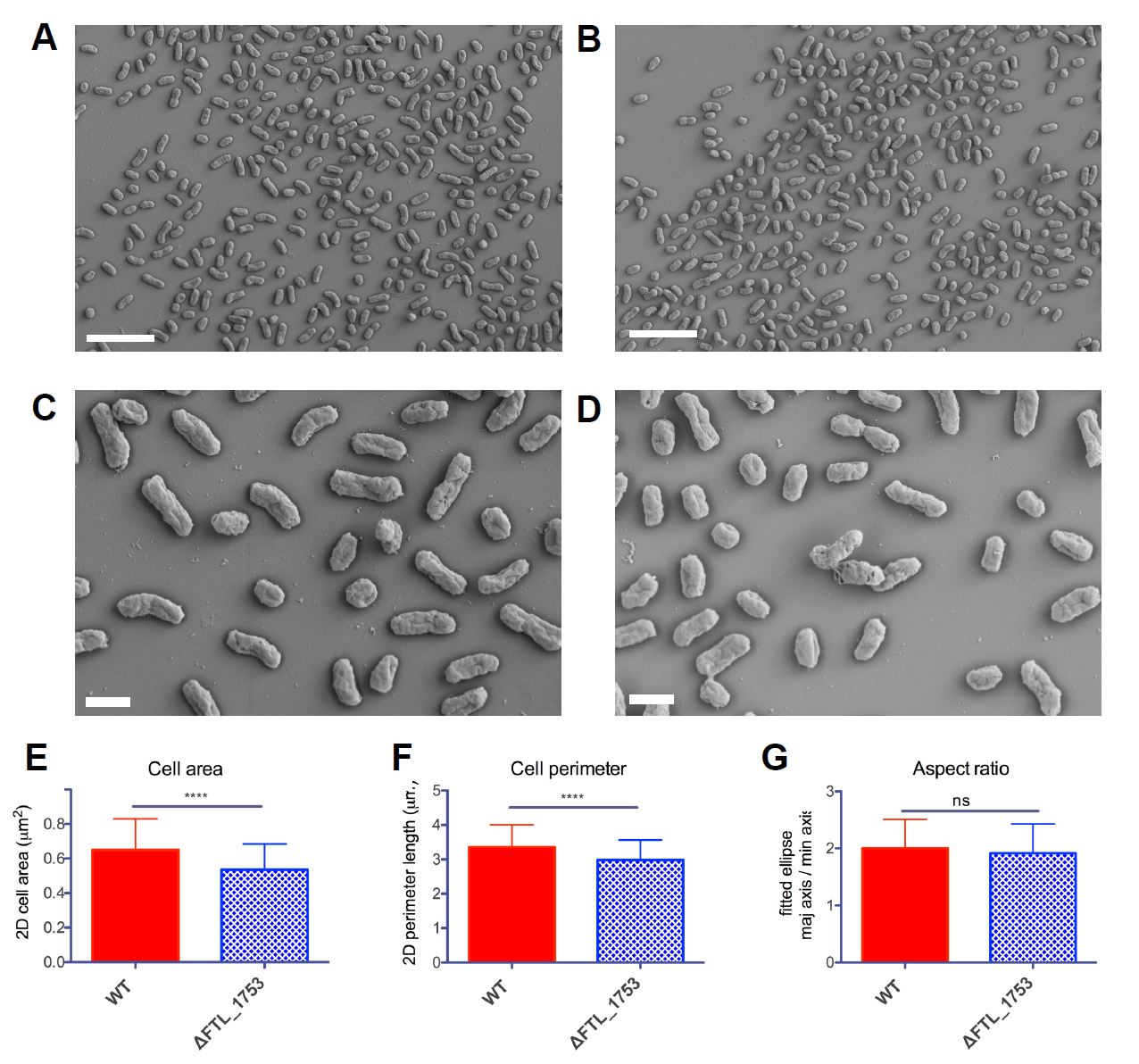
ΔFTL_1753 cells are smaller in average size compared to the wild-type. **A-B)** SEM images of wild-type (A) and ΔFTL_1753 (B) cells at 3,000X. Scale bars: 5µm. **C-D)** SEM images of wild-type (C) and ΔFTL_1753 (D) cells at 10,000X. Scale bars: 1µm. **E-G)** Cells (n>180 per group) were traced in Image J and subjected to the indicated analyses. E) 2-dimensional area of cell traces. F) the perimeter length of cell traces. G) the aspect ratio of ellipses fitted to cell traces. Two-tailed, unpaired t-test, ****p<0.0001. Negative staining experiments with uranyl acetate and phosphotungstic acid followed by TEM yielded similar results.

### Comparative bioinformatics reveal heterogeneity in the mechanosensitive channels encoded by F. tularensis subspecies

We chose to focus our study on the *holarctica* subspecies of *F. tularensis*, as this is the subspecies for which an aquatic cycle has been described, and which is responsible for outbreaks of tularemia from contaminated water. Nonetheless, as there is great interest in the highly virulent subsp. *tularensis* and also subsp. *novicida*, we compared the loci predicted to code for mechanosensitive channels in these subspecies to the loci present in subsp. *holarctica*. Of the four loci, the one corresponding to FTL_1753 was most highly conserved between the three subspecies, with only 1-2 amino acid substitutions between them (Fig. 5A and S5). Interestingly, the equivalent of FTL_1753 in the prototypical subsp. *tularensis* (Schu S4) genome is currently annotated as a pseudogene. However, our data and analyses suggest this is erroneous and should be corrected as explained in the discussion. The remaining three loci display marked heterogeneity between the *F. tularensis* subspecies. In the set of loci equivalent to FTL_0945, the C-terminal 44% of the protein is lacking in both ssp. *holarctica* and *tularensis* relative to ssp. *novicida* (Fig. 5B and S6). In another set, the N-terminal ~25% is absent in subsp. *holarctica* relative to the others (Fig. 5C and S7). In the final set, a frameshift has resulted in a premature stop codon and pseudogene annotation in subsp. *holarctica* (Fig. 5D and S8). Overall, these data suggest that: 1) the mechanosensitive channel found to protect *F. tularensis* subspecies *holarctica* from hypoosmotic shock in this study is conserved between the *F. tularensis* subspecies; and 2) the other loci predicted to code for mechanosensitive channels exhibit considerable heterogeneity between the subspecies.

**Figure 5:**
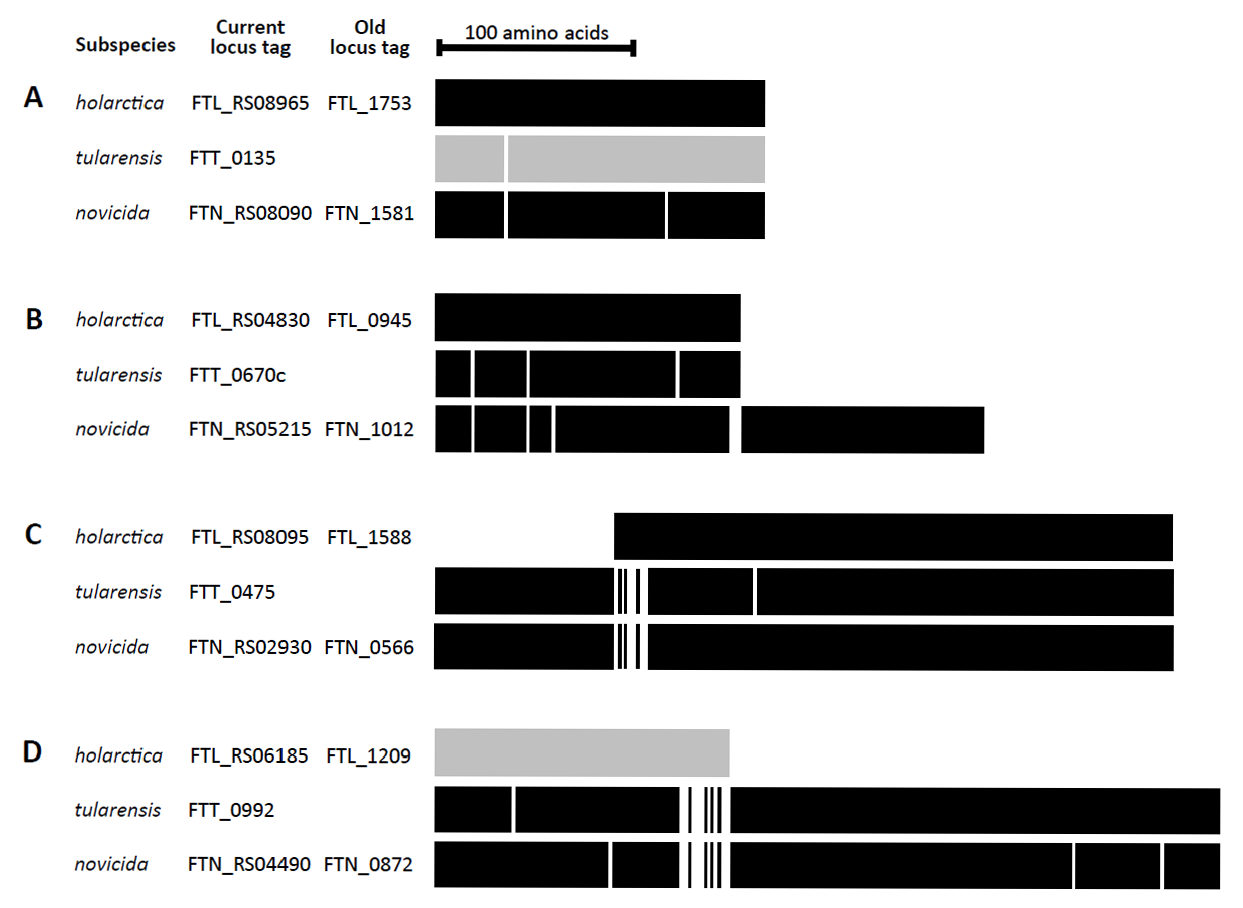
Summary of comparative alignments of loci predicted to encode mechanosensitive channels in *F. tularernsis* subspecies. White lines represent amino acid substitutions relative to subsp. *holarctica* except where no subsp. *holarctica* sequence is aligned. Gray shading signifies a current pseudogene annotation in RefSeq. For alignment detail, see supplementary Fig. 5-8.

## Discussion

The contamination of fresh water by *F. tularensis* subsp*. holarctica* is relevant to the bacterium’s ecological persistence and transmission to hosts, including humans. *F. tularensis* can only persist in fresh water for a limited time (27, 28), and is seasonally recovered from natural bodies of water, suggesting recurrent cycles of contamination (7, 31). Early soviet researchers reported that a single infected animal could contaminate 500,000L of water (44), and infected animals are indeed implicated in virtually all cases of water contamination with *F. tularensis* (6–8, 10–12, 16, 17, 19, 22, 26, 31–33). To survive an osmotic downshock, such as that encountered when going from a mammalian host to fresh water, bacteria rely on mechanosensitive channels which act as ‘pressure relief valves’. In this study, we have identified a mechanosensitive channel which protects *F. tularensis* subsp. *holarctica* from hypoosmotic shock (FTL_1753). The survival profile of ΔFTL_1753 upon downshocks of 100-300mM (Fig. 1B) closely resembles that of *E. coli* lacking both *mscL* and *mscS* (36), and is consistent with the notion that this is the single downshock-protective mechanosensitive channel in the subspecies. We demonstrated that the results of this study are applicable to clinical and environmental strains of subsp. *holarctica* (supplementary Fig. 1-3), and that bacteria lacking FTL_1753 are unable to survive the transition from the osmolarity of the mammalian host to fresh water (Fig. 1C and 3C). Notably, FTL_1753 has been found to be up-regulated within the intracellular growth niche(45); speculatively this may be to increase chances of survival in the event of a shift to a freshwater niche.

The presence of *F. tularensis* subsp. *holarctica* in fresh water has led to outbreaks of mostly oropharyngeal tularemia in at least 10 countries, as described earlier. Oropharyngeal tularemia is often initially misdiagnosed, and without appropriate treatment, it can become a chronic debilitating disease with the risk of serious complications (23). In addition, the aquatic cycle of subsp. *holarctica* can be implicated in many cases of ulceroglandular tularemia; water is thought to facilitate the spread of bacteria between animals and to mosquito vectors. In cases where humans develop ulceroglandular tularemia from mosquito bites, mosquitos may acquire the bacterium from water during their development. In Sweden, where most cases of tularemia are associated with mosquito bites (3, 46), a strong association has been found between infection and recreational areas near water (47). Furthermore, mosquito larvae reared in pond water from endemic areas, or experimentally exposed to *F. tularensis* during development, have been shown to harbor *F. tularensis* as adults (48, 49). In cases where humans develop ulceroglandular tularemia from handling infected animals, *F. tularensis* can often be found in water within the animals’ habitat (26, 33, 50); the presence of viable bacteria in the water is thought to contribute to the spread and persistence of infection among animal populations (26, 33). Thus, the mechanosensitive channel identified here is likely to be important to the overall ecological persistence of *F. tularensis* and its transmission to humans and other susceptible hosts.

Bioinformatically, we identified four loci predicted to contain mechanosensitive channel domains in each *F. tularensis* genome surveyed, including in subsp. *tularensis* and subsp. *novicida* (Fig. 5, S1-S3, S5-S8). Notably, FTL_1753 was the most conserved between the subspecies, with only one amino acid substitution between ssp. *holarctica* and *tularensis* (Fig.5A, S5). At 165 amino acids, FTL_1753 is in the smallest 2% of the mechanosensitive channel superfamily, which contains over 69,000 members. It is also considerably smaller than MscS in the model organisms in *E. coli* and *B. subtilis*, which are approximately 280 amino acids. This has led to a ‘partial’ annotation in the non-redundant protein record for FTL_1753 (WP_003017262.1), and a pseudogene annotation in the manually curated subsp. *tularensis* SchuS4 genome. Our data demonstrate that despite its small size, this gene does indeed code for an active mechanosensitive channel that protects from downshock. The other loci containing mechanosensitive channel domains exhibited considerable heterogeneity between subspecies. The functions of these remain to be determined, but in subsp. *holarctica* may include roles other than osmotic shock protection. For instance, other members of the superfamily have been shown to play roles in signal transduction (51).

In conclusion, we have identified a small, 165 amino acid mechanosensitive channel that protects *F. tularensis* subsp. *holarctia* from downshock. This channel is required for the bacterium to survive the transition from mammalian hosts to fresh water, and likely contributes to ecological persistence of *F. tularensis* and transmission between hosts.

## Materials and Methods

### Bioinformatic analysis of msc-domain containing genes in F. tularensis

The amino acid sequences of mechanosensitive channels (MscS & MscL) that protect *E. coli* from downshock (36) were retrieved from UniprotKB, and input into the Conserved Domain Architecture Retrieval Tool (CDART) (37). Results were filtered to the *Francisella* genus using the NCBI taxonomy tree to identify genes containing msc domains in the strains of interest. ORF and gene sequences were retrieved from RefSeq genomes, compared using standard protein BLAST and Needleman-Wunsch global alignment, and visualized using Multiple Align Show (http://www.bioinformatics.org/sms/multi_align.html).

### Bacterial strains and growth conditions

*Francisella tularensis* subsp. *holarctica* Live Vaccine Strain (LVS) was obtained from Albany Medical College and was grown on MH chocolate agar (Mueller-Hinton II agar (BD BBL) supplemented with 1% hemoglobin (remel) and 1% isovitalex (BD BBL)). Liquid cultures of *F. tularensis* were routinely grown in Brain Heart Infusion Broth, Modified (BHI, BD BBL #299070) at pH 6.4-6.8. Cloning procedures were performed using DH5α *E. coli* grown in LB broth, Miller (BD Difco) or LB agar, Miller (EMD chemicals). Liquid cultures were incubated at 37°C in an orbital shaker operating at 200rpm. Agar plates were incubated at 37°C, 5% CO_2_ for 48-72 hours.

### Primers, plasmids and DNA manipulation

All primers used in this study are shown in table 1. The pMP812 *sacB* suicide vector (38) was used to generate targeted, marker-less, in-frame deletions of desired genes. Briefly, 700-900 base pair PCR products flanking the sequence to be removed were amplified, digested and ligated together, then re-amplified as one unit and cloned into the mcs of pMP812. A complementation plasmid for *mscS* was generated by modifying the pKK214-GFP plasmid (29). First, the GroEL promoter and GFP gene were removed by digestion with *Xba*I and *Eco*RI. A multiple cloning site with four restriction sites was then added by annealed oligo cloning, with phosphorylated MCS_4__1 & _2 oligonucleotides, to generate pKK214-MCS_4_. The bacterioferritin promoter and FTL_1753 were amplified, digested and ligated together, then re-amplified as one unit and cloned into pKK214-MCS_4_ with EcoRI. The resulting plasmid is referred to as p-1753. All oligonucleotides were synthesized by Integrated DNA Technologies or the Penn State Genomics Core Facility. All enzymes were from New England Biolabs. Vent_R_ DNA polymerase was used for cloning and Taq DNA polymerase for screening. Genomic DNA was isolated from *F. tularensis* using an E.Z.N.A. Bacterial DNA Kit (Omega Bio-Tek) and plasmid DNA was isolated using an E.Z.N.A Plasmid DNA Mini Kit I (Omega Bio Tek).

**Table 1:**
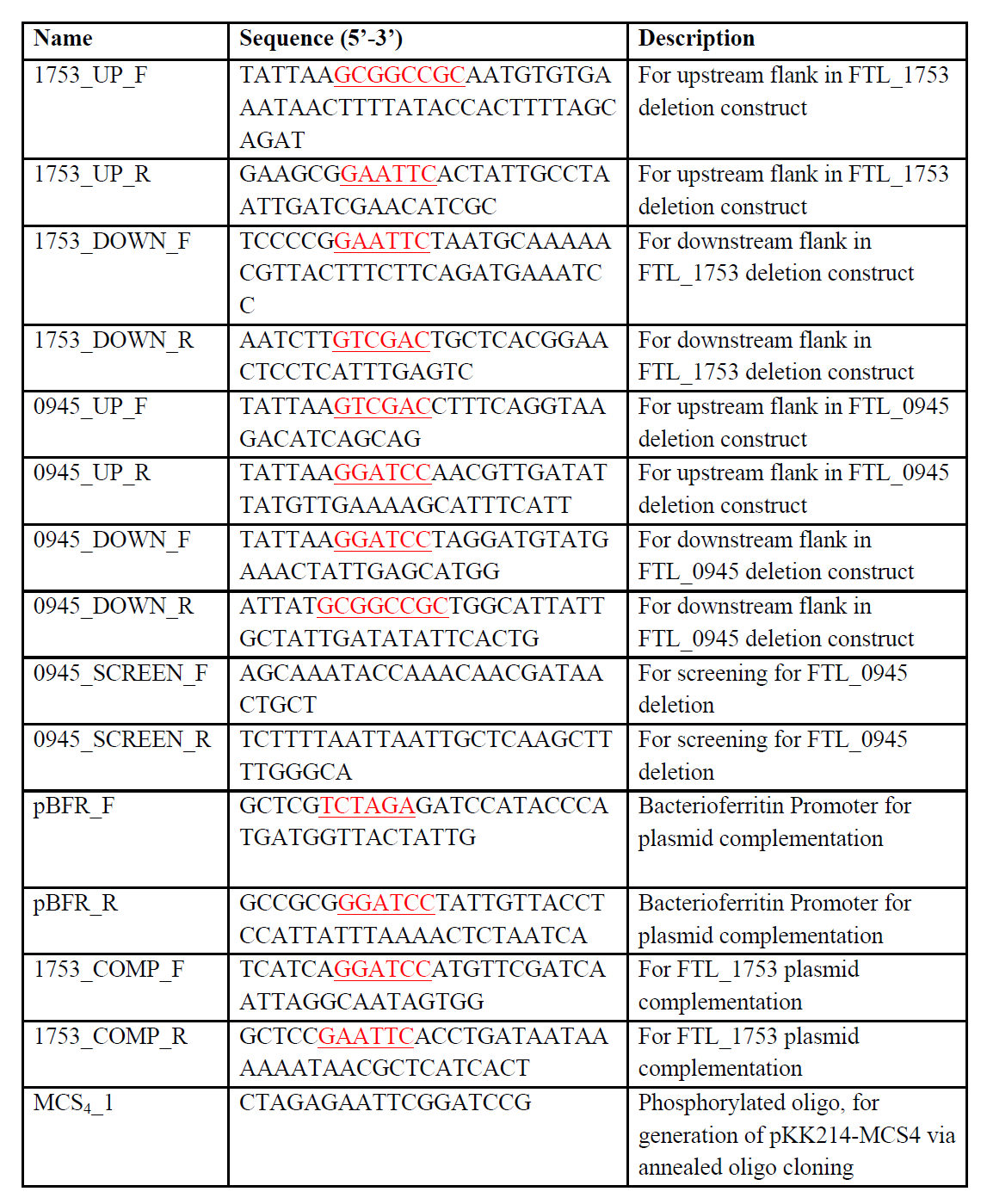
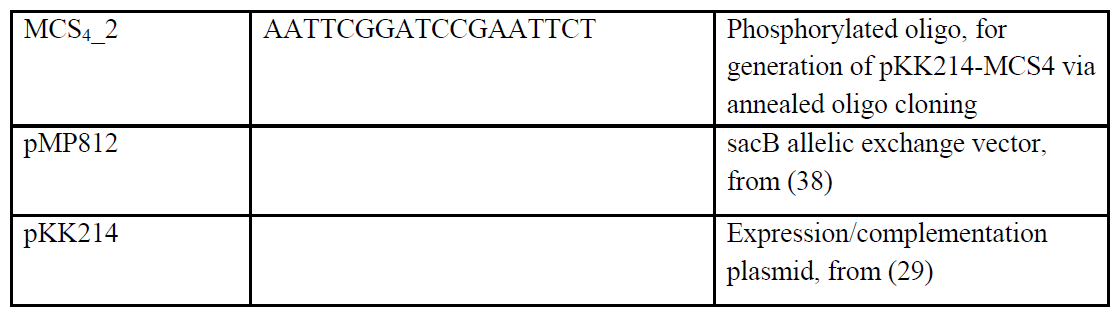
Primers 603 and plasmids used in this study

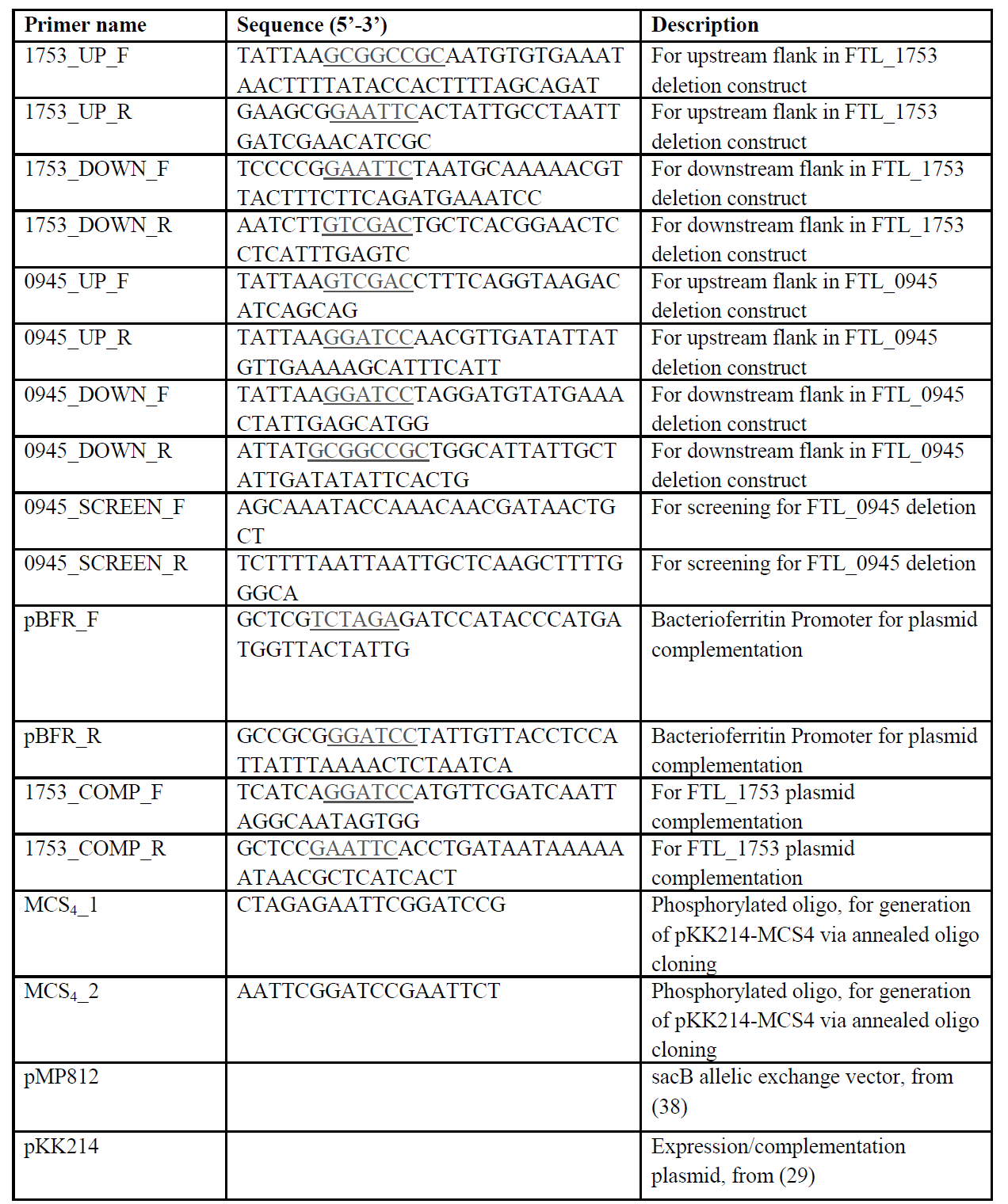

### Generation of in-frame deletions via allelic exchange and trans-complementation

Marker-less, in-frame deletions of FTL_1753 (new locus tag FTL_RS08965) and FTL_0945 (new locus tag FTL_RS04830) were generated using the pMP812 *sacB* suicide vectors as previously described (38). PCR was used to screen for secondary recombinants with deletions, as outlined in Fig. S4. *Trans*-complementation of FTL_1753 was achieved by introducing the p-1753 plasmid into a confirmed ΔFTL_1753 strain. Electrocompetent *F. tularensis* cells were prepared as previously described (39), except that electroporations were done using 0.1cm gap cuvettes (VWR) and Bio-Rad micropulser using Ec1 settings (1.8kV, 10µF and 600Ω).

### Hypoosmotic shock assays

For downshock experiments conducted in media, single colonies of each strain were picked from MH-chocolate agar plates and grown in BHI supplemented with 300mM NaCl (EMD chemicals). Mid-log cultures were diluted (in BHI +300mM NaCl) to a concentration of approximately 5x10^5^ CFU/mL, then 20µL of each culture added to 980µL of BHI +300, +200, +100 or +0 mM NaCl, to produce hypoosmotic shocks of 0, 100, 200 and 300mM, respectively. The resulting cultures were vortexed and incubated for 1hr at room temperature, before plating on MH chocolate agar with the same concentrations of NaCl added to prevent further osmotic shock. Percent survival was calculated by comparing viable counts from cultures subject to downshock to cultures maintained with 300mM NaCl throughout the experiment. For downshock experiments from PBS to fresh water, a similar method was used, except that no NaCl was added to growth media, and cells were equilibrated in PBS for 30 minutes before dilution into fresh water for 10 minutes. To test a source of water more naturally relevant than double-distilled H_2_O, water was retrieved from a freshwater lake in Pennsylvania and passed through a 0.2µm filter.

### H_2_O_2_ and detergent sensitivity assays

Single colonies of each strain were picked from MH-chocolate agar plates and grown in BHI. Early-log phase cultures were diluted to a concentration of approximately 4x10^4^ CFU/mL, and added 1:1 to plain media or media with 2X the final desired concentration of detergent or H_2_O_2_. All dilutions and challenges were performed in identical media, such that the only variable would be the detergent/ H_2_O_2_. These cultures were vortexed and incubated for 1 hour at room temperature, before being diluted 1/10 in BHI, and 100µL plated to enumerate viable CFUs. Percent survival was calculated relative to the negative control plates for the same strain (which yielded ~200 CFU).

### Determination of antibiotic minimum inhibitory concentrations

2-fold serial dilutions of each antibiotic were prepared and added to 96 well plates. Bacterial cultures were grown overnight, diluted to an OD_600_ of 0.050, and added 1:1 to plate wells containing media/antibiotics, in triplicate. Plates were incubated overnight (~18h), and bacterial growth determined by measuring optical density at 600nm. The MIC was defined as the lowest final concentration of antibiotic for which cultures did not grow beyond an OD_600_ of 0.1.

### Intra-macrophage replication and macrophage-to-freshwater downshock assays

Raw 264.7 macrophages were maintained in complete DMEM (DMEM +10% FBS, 1mM sodium pyruvate, 2mM L-glutamine, 10mM HEPES, 1X non-essential amino acids). For intra-macrophage replication assays, Raw 264.7 macrophages were seeded at a concentration of 3x10^5^ cells per well in 24 well tissue culture plates (Greiner Bio One) the evening before the assay. Macrophages were infected with early-log, BHI-grown *F. tularensis* at a multiplicity of infection of 100 bacteria per macrophage. Plates were centrifuged to facilitate synchronized infection (10 minutes, 300 *x g*, RT). After a one hour incubation to allow for bacterial uptake, media was aspirated and replaced with media containing 100µg/mL gentamicin (Gibco) to kill extracellular bacteria. Following an additional one hour incubation, cells were washed twice with PBS. For wells designated for later time points, fresh media (without antibiotic) was added. Cells in other wells were lysed by the addition of 100µL/well of 0.1% w/v sodium deoxycholate (sigma) in PBS. After lysis was observed, 900µL of PBS was added to each well, and serial dilutions plated on MH chocolate agar to determine the number of bacteria that had been taken up by macrophages. 20 hours post-infection, cells were similarly lysed and plated to determine the number of bacteria present. Macrophage-to-freshwater downshock assays were done in a similar fashion but with the following alterations: 5x10^4^ cells were seeded per well to minimize the contribution of intracellular salt carryover during lysis, and cells were lysed at the 2 hour timepoint only using 200µL/well of either ddH_2_O or PBS +0.1% w/v sodium deoxycholate.

### Mouse infections

C57BL/6J mice were maintained in specific pathogen-free conditions at the Pennsylvania State University animal care facilities. Isoflurane-anesthetized mice (n=8 per group) were infected intranasally with 10,000 CFU of bacteria in 50µL of PBS. Serial dilutions of the inoculum were plated on MH chocolate agar to confirm the correct dose was administered. Body weight was measured daily, and mice that lost 20% or more of their starting body weight were euthanized and counted as having succumbed to infection, per Institutional Animal Care and Use Committee (IACUC) guidelines. Experimental groups were age and sex matched.

### Electron Microscopy

5mL bacterial cultures were grown to an OD_600_ of approximately 0.5 and rinsed twice by centrifuging at 1500 x *g* for 4 minutes then resuspending in pre-warmed media. Cells were fixed by resuspending in 2% glutaraldehyde (Electron Microscopy Sciences) in PBS for 30 minutes at room temperature, then overnight at 4°C. The cell suspension was applied onto Poly-L-Lysine coated coverslips for 10 min at 4°C and proceeded for critical point drying in a chamber of Leica EM CPD300 (Leica Microsystems, Buffalo Grove, IL) critical point dryer through an automatic program. The coverslips were then mounted onto a double-sided carbon tape on a single pin aluminum SEM stab (Ted Pella, Redding, CA) and sputter coated with iridium on a rotation stage of Leica ACE600 (Leica Microsystems, Buffalo Grove, IL). The bacterial samples were examined and imaged in a Zeiss Sigma FE-SEM (Carl Zeiss Microscopy, Thornwood, NY) at 2 kV. The images were analyzed to determine cell 2D area, perimeter length, and the aspect ratio of a fitted ellipse using ImageJ v1.50i(40).

### Statistics

Statistical differences were determined as indicated in figure legends using GraphPad Prism 5.0.

### Ethics statement

All animal experiments were carried out by following recommendations and approval from the Pennsylvania State University Animal Care and Use Committee (protocol 46070) with great care taken to minimize suffering of animals.

## Acknowledgements

We sincerely thank Dr. Martin Pavelka, University of Rochester, For providing the pMP812 plasmid, and Dr. Karl Klose, University of Texas at San Antonio, for providing the pKK214-GFP plasmid. We also thank The Huck Institutes of the Life Sciences for the facilities and technical support.

## Funding Information

This study was supported by a T32 training grant AI 074551 (NIAID/NIH) to DRW, AI077917 (NIAID/NIH), AES4605 (USDA) and start up finds from Penn State University and The Huck Institutes of the Life Sciences to GSK. The funders had no role in study design, data collection and interpretation, or the decision to submit the work for publication.

